# Phenotypic changes of bacteria through opportunity and global methylation leads to antibiotic resistance

**DOI:** 10.1101/2020.05.27.114090

**Authors:** Muniasamy Neerathilingam, Sumukh Mysore, Sneha Bairy, Chetan Chandola, Venkadeshwari Varadharajan, Ram Rajasekharan

## Abstract

The antibiotic stress on bacteria leads to initiation of adaptive mechanisms, including exploiting the available opportunity, if any, for cell survival. In order to use the opportunity for survival while under threat, the microbe undergoes various mechanisms which are not completely known e.g. homologous recombination, horizontal gene transfer etc. Our aim is to understand the adaptive mechanism for cell survival during stress, especially antibiotic stress, in *E. coli* in the presence of opportunity. Understanding this mechanism in bacteria that gained resistance will help in identifying alternative survival pathways. By subjecting a recombination deficient (ΔRecA) strain of bacteria to antibiotic stress, we expected cell death, due to its inability to repair DNA damage (*1, 2*). Here we show that providing an opportunity in the form of an antibiotic resistance gene with homologous ends aids bacterial survival. There was 3-fold increase in cell envelope thickness along with 2.5-fold increase in phosphatidylethanolamine (PE) content, and enhanced antibiotic resistance to >4000µg/mL (Kan). We observed genome-wide alteration of methylation pattern that lead to changes in transcriptome, proteome, lipidome, and metabolite level, thus, leading to morphological and physiological changes. We prove that global methylation helps in survival of bacteria under stress that changes essential pathways like energy, cell envelope, lipids, amino acids acid, etc. leading to over production of cell wall components including synthesis of PE. By inhibiting the activity of methyltransferase, it is noticed that there is reduction in PE synthesis in agreement with demethylation. This proves that the phenotypic changes are caused due to the global methylation, and also demonstrates that demethylation could be used as a strategy to prevent antibiotic resistance in microbes.

**One Sentence Summary:** Global methylation determines the survival of bacteria to gain the antimicrobial resistance with an opportunity

---

Antibiotic stress in bacteria is overcome by various adaptive mechanisms, which are opportunistic (*3*). The antibiotic stress on bacteria makes it struggle for its survival and is overcome by various adaptive mechanisms, including exploiting the available opportunity, if any. In order to use the opportunity for survival while under threat, the microbes undergo certain mechanisms which are not completely known. For example, how it is primed for withstanding the stress condition including the presence of antibiotics, prior to using the opportunity and upon exploiting the opportunity of how it adapts to the environment. The understanding of this adaptive mechanism for cell survival is also fundamental to deal with antimicrobial resistance (AMR). Bacteria especially hide from antibiotics by changing its cell wall thus avoiding stop penetrating into bacteria.

To understand the effect of antibiotic stress and the adaptations undergone by the organism for its survival, homologous recombination is one of the many mechanisms necessary for overcoming external stress in prokaryotes and eukaryotes, particularly during a DNA double-strand break on the bacterial genome(*4*). When the event occurs in bacteria, it acts as a survival mechanism and causes its evolution into drug-resistant cells, which may make the infection untreatable. Hence, it is paramount to study the effect of antibiotic stress and the adaptations undergone by the organism for survival. In order to understand the phenomena, we used the strain of *E. coli* (ΔRecA) in which the RecA enzyme has been removed (Supplementary table S1), thus allowing us to distinctly study the effects of stress (antibiotic) induced recombination events in the organism. We used homologous recombination model to mimic such an environment. The possibilities of occurrence of other homologous recombination RecFOR and RecBCD events are nullified upon removal of RecA (*5-8*) and RecE pathway is non-functional without an additional sbcA mutation(*8, 9*). However, the recombination event in the chosen strain (ΔRecA) is possibly through RecA independent pathway(*10*). We introduced antibiotic as external stress to the bacteria, and the subsequent survival of bacteria would depend on the opportunity provided. Herein, the opportunity is the formation of a circular plasmid through recombination of four fragments (4FR). The four fragments with homologous sequence details are provided in Fig. 1a and in the supplementary table S2 with the antibiotic resistance gene (Kan^r^) and ColE1 (ori).

**Fig. 1a.**
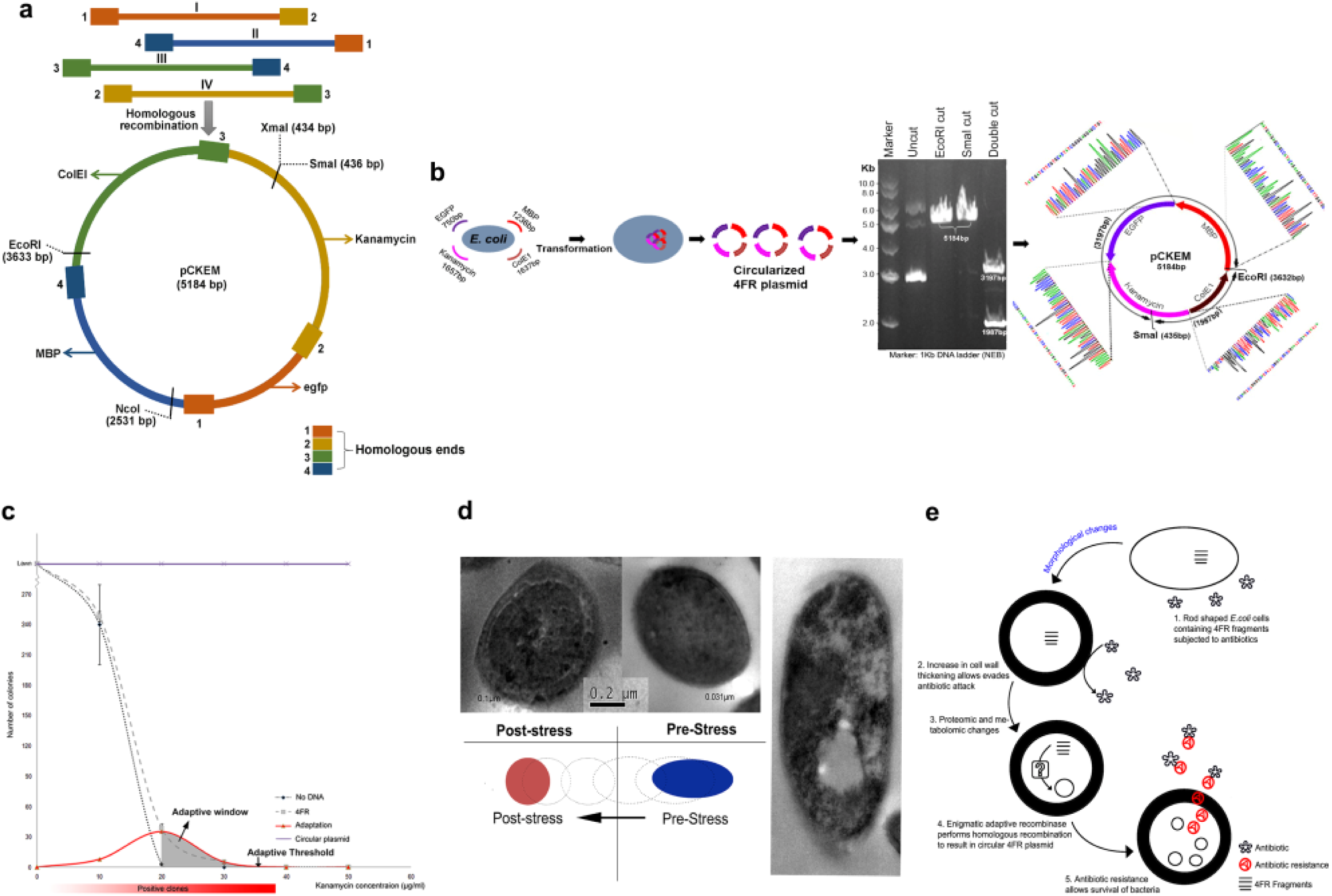
Design of linear constructs of four fragments (4FR) for the formation of a plasmid (pCKeM) by homologous recombination (1, 2, 3 and 4 corresponds to homologous ends). All the individual fragments have been designed with a homologous sequence at both 5’and 3’end for recombination. Fragment I codes for an enhanced green fluorescent protein (egfp) with 1 and 2 ends contains a homologous sequence of fragment II and IV. In case of Fragment II which contains Maltose Binding Protein (MBP) with two ends of a homologous sequence of fragment III and I. 5’and 3’ends of Fragment III are homologous to Fragment IV and II along with it carries the origin of replication colicin-E1 (ColEI). The other fragment, IV consists of a kanamycin resistance gene has a homologous sequence to I and III ends. All the four fragments I, II, III and IV and the pCKeM map are shown in Figure (1a). Sources for amplifying each fragment and primers used for generating the linear fragments with its amplicon size are detailed in Supplementary Table S2. The complete sequence details for all the four fragments are given in Supplementary file S1. This comprises homologous sequences highlighted with different colors. **1b.** The procedure adapted for the creation of pCKeM by homologous recombination using 4FR transforming into the *E. coli* BLR strain. Confirmation of the generated pCKeM was determined by both single and double restriction and digestion using *SmaI* and *EcoR*I which provides the expected size of 5184 bp for single digestion and 3197 bp and 1987 bp for double digestion. The occurrence of homologous recombination for pCKeM was confirmed precisely by using the Sanger sequencing method. **1c.** An opportunity has driven extension of survival by 4FR formation within the adaptive threshold and its adaptive window. ΔRecA strain containing kanamycin-resistant gene harboring plasmid pET28a (no stress) and transformed 4FR into ΔRecA strain were plated from sub-inhibitory to lethal dosages of kanamycin as shown in the figure. Untransformed ΔRecA strain was kept as a control to observe the emergence of sub-inhibitory colonies. A number of 4FR colonies (pCKeM) were plotted against the antibiotic concentration in triplicates. The adaptive curve was obtained by subtracting the number of colonies obtained in 4FR with the values of the sub-inhibitory curve and the adaptive window drawn in a red curve (the area shaded with grey) which describes the extension of survival by 4FR formation (pCKeM). The adaptive threshold is determined where no colonies were found as marked in the figure. **1d.** Pre and Post stress showing a difference in cell envelope thickness which might facilitate the stress-induced adaptive recombination that results in kanamycin resistance of 4FR recombination to form pCKeM. The post-stress of ΔRecA strain was transformed with 4FR (four fragments). The pre-stress of ΔRecA strain was transformed with pCKeM which was isolated from the post-stress-strain. **1e.** Schematic representation of Fig. 1d. for cell envelop thickening the occurrence of adaptive recombination.

We observed that the independent homologous recombination of four fragments (4FR) to form circular plasmid (Fig. 1a) in ΔRecA strain takes place only upon exposure to the antibiotic stress that was confirmed by performing two different adaptive test experiments as described in the Extended Data Fig. S1a, b. The recombined 4FR plasmid (pCKEM) from the ΔRecA strain was later sequenced (Supplementary File S1) to confirm the occurrence of the homologous recombination. We verified its size by restriction and digestion (Fig. 1b).

We observed an adaptive window of the antibiotic concentration of 20-30 μg/ml within which the four-fragment recombination (4FR) took place (Fig. 1c). Below a concentration of 20 μg/ml, the organism survived without undergoing recombination, as the stress was too low. Whereas, more than a concentration of 30 μg/ml, the organism was unable to carry out recombination due to high stress. Providing the opportunity of 4FR to the cells ameliorates the organism to withstand a sub-inhibitory dosage from 20 to 30 μg/ml. A tolerance has previously been attained in *E. coli* K12, and our control by treatment with sub-inhibitory dosage (20 μg/ml) of kanamycin shows that the organism cannot overcome the antibiotic stress beyond the lethal dosage (20 μg/ml). The control strain is *E. coli* BLR transformed with the isolated circular plasmid (pCKEM) formed by four-fragment recombination. Here, upon plotting the subtraction of the sub-inhibitory curve and 4FR values, a bell curve is obtained (Fig. 1c). This bell curve revealed the window created by the first few 4FR clones, in which this 4FR joining was observed.

To better understand the phenomenon, the morphology of the control and 4FR strains were analyzed using TEM; we observed that there was a three-fold increased thickness in the cell-envelope of the organism (4FR) as compared to the control (Fig. 1d). The thickened cell envelope could have prevented entry of the kanamycin, thereby providing resistance as depicted in Fig. 1e. The control and 4FR were plated on different concentration of kanamycin (250, 500, 1000, 2000 and 4000 μg/ml). The 4FR is resistant to more than 4000 μg/ml while control was up to 500 μg/ml only (Extended Data Table S1).

To understand the changes in cell envelope thickening and the high antibiotic resistance gained in 4FR, we isolated total lipids from the control and 4FR and separated by 1D and 2D TLC (Fig. 2a). The total lipids were used for the lipid profile analysis. It was observed that there was a 2.5 fold increase in phosphatidylethanolamine (PE) content in 4FR as compared to control (Fig. 2b). The lipidome was analyzed by injecting the samples into the AB Sciex TripleTOF™ 5600 System using the MS/MS ALL method. Further, we observed no significant difference in fatty acid composition (Fig. 2c, d). Since the major component in the cell envelope of gram-negative bacteria is PE, we suspected that the increased thickness of the cell envelope that is observed in the TEM could be due to PE.

**Fig. 2a.**
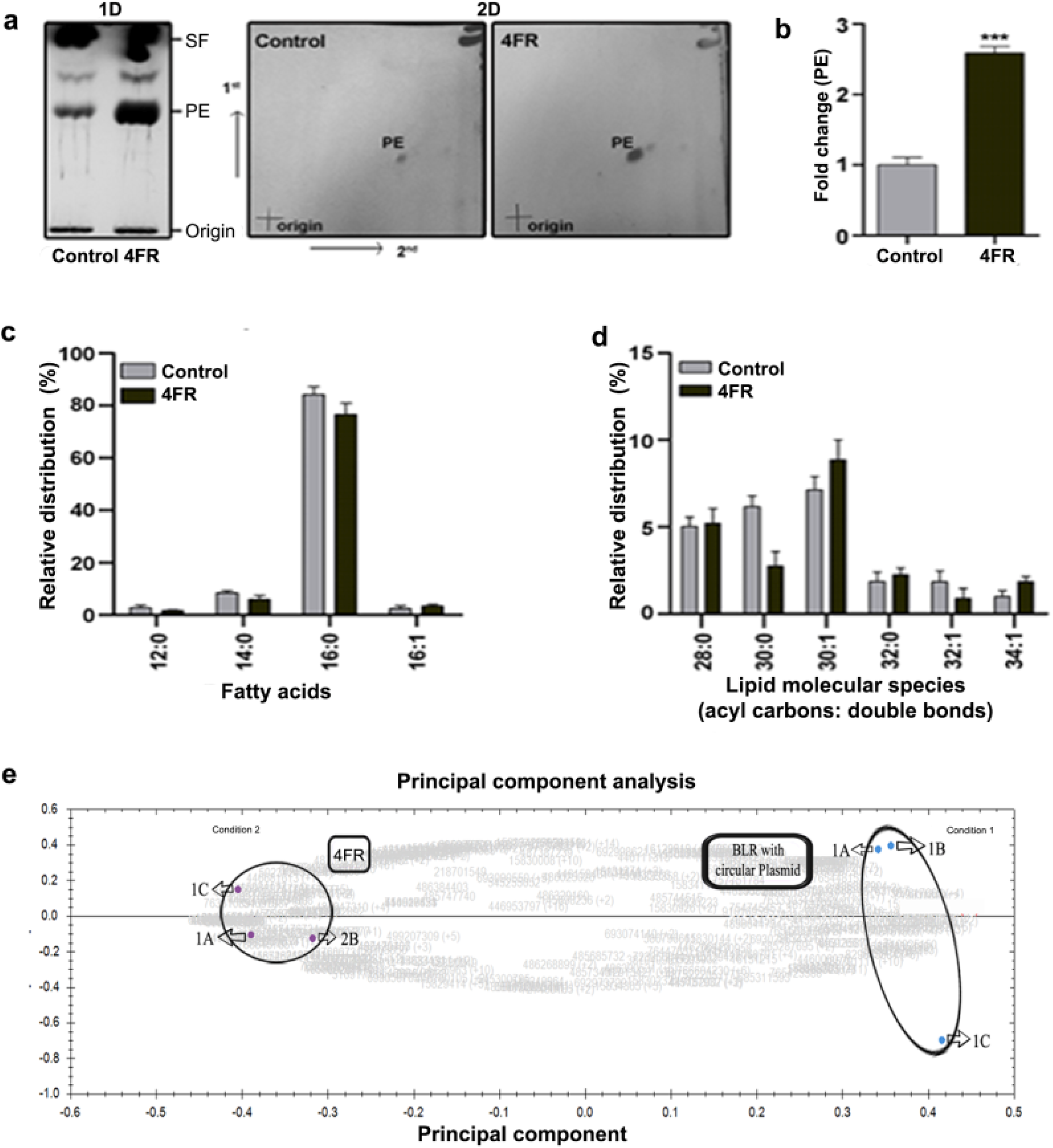
Phosphatidylethanolamine content and molecular species in 4FR, Left panel, one-dimensional-silica TLC analysis. The cells were harvested, and extracted lipids were resolved on a TLC plate. PE, phosphatidylethanolamine; SF, solvent front. Right panel, two-dimensional-TLC analysis. The lipids were separated by using chloroform: methanol: ammonia (65:25:5, v/v) as the first-dimensional solvent system and chloroform: methanol: acetone: acetic acid: water (50:10:20:15:5, v/v) as the second-dimensional solvent system. Lipids were identified by charring the plate with copper sulfate. **2b.** A relative fold change of PE in 4FR. The values are represented as the mean (±SD) of three independent experiments performed in triplicates. Significance was determined at *p<0.001. **2c, d.** Relative distribution of fatty acids and molecular species. The lipids were extracted and analyzed by MS/MS^ALL^ (three independent experiments performed in triplicates). **2e.** 2D PCA analysis of the proteomics data obtained using label-free LC-MS showing distinct separation between the 4FR and control. Data were acquired using Thermoscientific LTQ Orbitrap Discovery. Further, proteomic data were filtered and analyzed in Progenesis QI blasted across swiss-prot and RefSeq databases at a p-value of 0.05%.

In order to understand the high-level PE synthesis and envelope thickness, we employed to study the proteomic profile of both 4FR and control using label-free LC-MS. There is a distinct separation between 4FR and control observed through 2D PCA (Fig. 2e). It shows a noticeable difference of 71 proteins, in that 52 are identified and involved in the carbohydrate metabolism, stress-related, nucleotide metabolism, cell wall synthesis, and amino acids metabolism and the rest are hypothetical proteins (Extended Data Table S2). The estimation of amino acids and few other metabolites using UHPLC-MS/SRM method(*11*) (Fig. 3a) shows the correlation for the changes in the metabolism and its pathway corresponding to proteomic data (Fig. 3b). Some of the differently regulated proteins show significant changes in the expression. For example, ribosomal protein rpsE that is involved in conferring antibiotic resistance which was one of the significant protein is found to be up-regulated(*12*).

**Fig. 3a.**
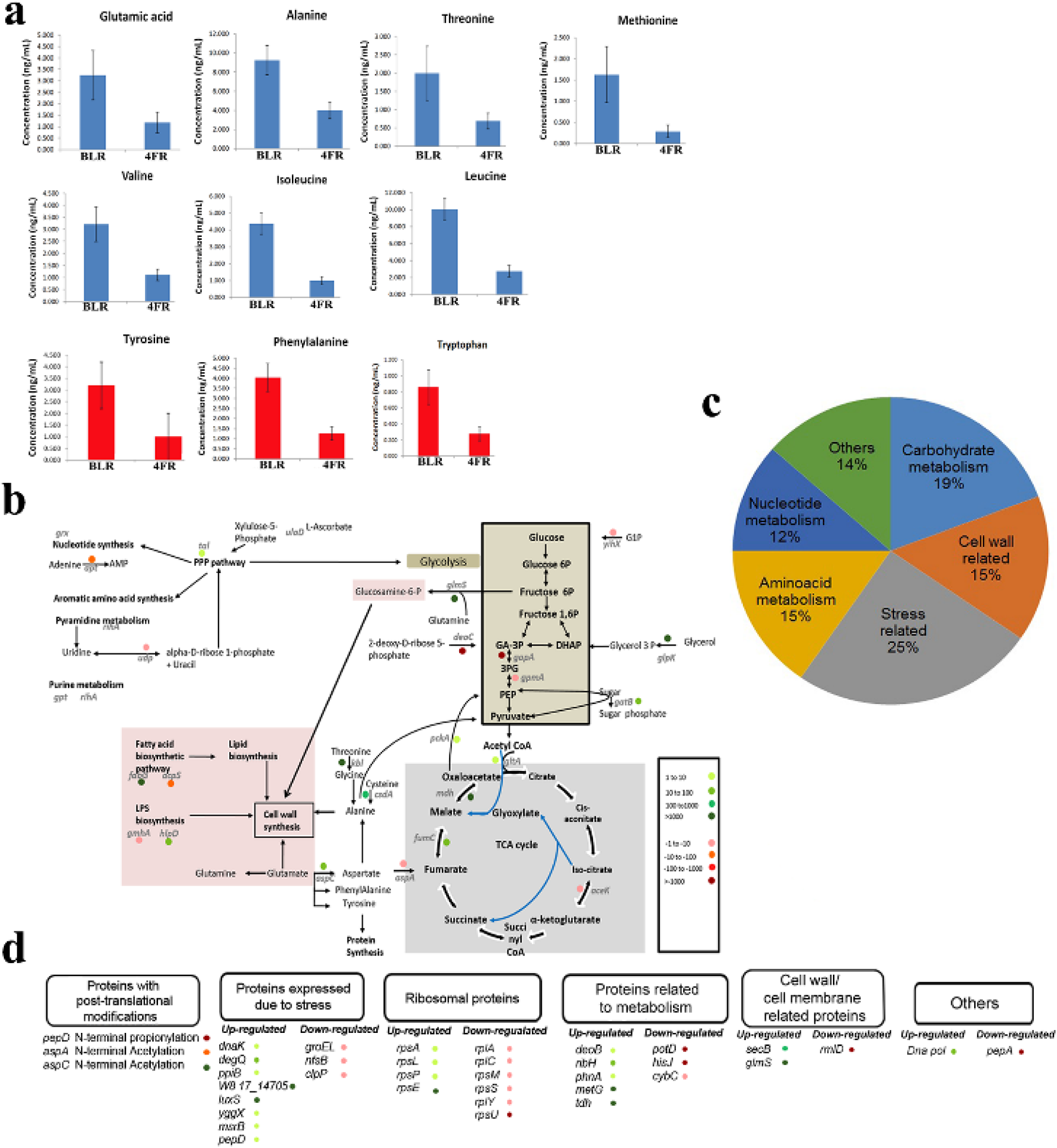
Non-aromatic amino acids (blue chart) and aromatic amino acids (red chart) showing a significant difference between control vs 4FR strains pre (BLR) and post-stress (4FR) strains were compared. **3b.** Illustration of comparing pre and post-stress in the RecA strains analyzed by Label-free quantification. The differentially regulated enzymes were first listed and then fit into glycolysis, TCA cycles, cell wall synthesis, and PPP pathways. These pathways are all interlinked by certain substrates/products and the holistic view gives us a better understanding of the overall post-stress effect in 4FR. Here, proteins up-regulated in 4FR are shown in shades of green and protein down-regulated in 4FR are shown in shades of red. **3c.** Distribution of differentially regulated genes in the 4FR clone represented in the form of a pie chart. It shows that a significant percentage of the genes are involved in cell wall synthesis, metabolism, and stress. **3d.** The enzymes that were differentially regulated but not fit into the pathway in Fig 3b are categorized and listed out under different sub-headings.

The analysis of the metabolic pathways shows the downregulation of enzymes involved in glycolysis (Fig. 3b), which could be compensated by the up-regulation of the enzymes involved in glycolysis feeder pathways such as glycerol kinase (*13*). An up-regulation in the glyoxylate pathway is indicated by the up-regulation of enzymes common to both glyoxylate and TCA cycle and downregulation of isocitrate dehydrogenase, which is unique to the TCA cycle. It could be because of the insufficient transport of glucose inside the cell, which triggers the glyoxylate pathway to assimilate C4 compounds that can be used for gluconeogenesis(*14*). Alanine and glutamate were observed to be at lower levels in the 4FR clone (Fig. 3a). We observe an over-expression of enzymes involved in alanine synthesis in at least two different pathways (csdA and kbl). Kbl is an enzyme that converts threonine to glycine, that in-turn can be converted to alanine(*15*). This explains the lower levels of threonine in the 4FR clone. The downregulation of the enzyme aspA, which converts aspartate to fumarate, allows for the conversion of aspartate to alanine. Further, the decrease in alanine content in the 4FR (Fig. 3a) indicates that alanine could be recruited for cell wall synthesis by the formation of a Pentapeptide l-Ala-γ-d-Glu-l-lysine-d-Ala-d-Ala binding to the MurNAc group(*16*). The proteomics data showed an over-expression of glmS, a protein responsible for the production of glucosamine-6-phosphate, which is a sugar moiety present in the cell wall as NAG (N-Acetyl Glucosamine). There is also a highly over-expressed fabG, involved in the fatty acid biosynthetic pathway, which in-turn might be involved in lipid biosynthesis required for the synthesis of thicker outer membrane synthesis(*17*). rmlD, which is greatly down-regulated in the 4FR clone is an enzyme involved in LPS bio-genesis(*18*). A drastic decrease in the methionine content and up-regulation of tRNA methionine synthetase, involved in covalently linking methionine with its cognate tRNA in 4FR indicates that the methionine has been utilized for protein synthesis (Fig. 3a). The low content of aromatic amino acids, such as phenylalanine, tyrosine, and tryptophan, in the 4FR, could also direct towards protein synthesis (Fig. 3a).

To address the changes, we performed a whole-genome sequence for 4FR and control using long-read sequencing with the MinION (R7.3). We observed that there was no change in the nucleotide sequences of the genome of both strains (Fig. 4a). In this study, we applied a probabilistic strategy that helps the expansion of the nucleotide alphabet to include bases containing chemical modifications and perform a detailed methylation analysis. Upon confirmation of methylation in the hot spot genes in 4FR genome reveals the precision of this method (Extended Data Table S4a). In addition, the tetracycline resistance gene (tet) in the 4FR which has been already inserted in the genome found to be methylated (Extended Data Table S4b), hence 4FR sensitivity to tetracycline while control is resistant to the tetracycline.

**Fig. 4a.**
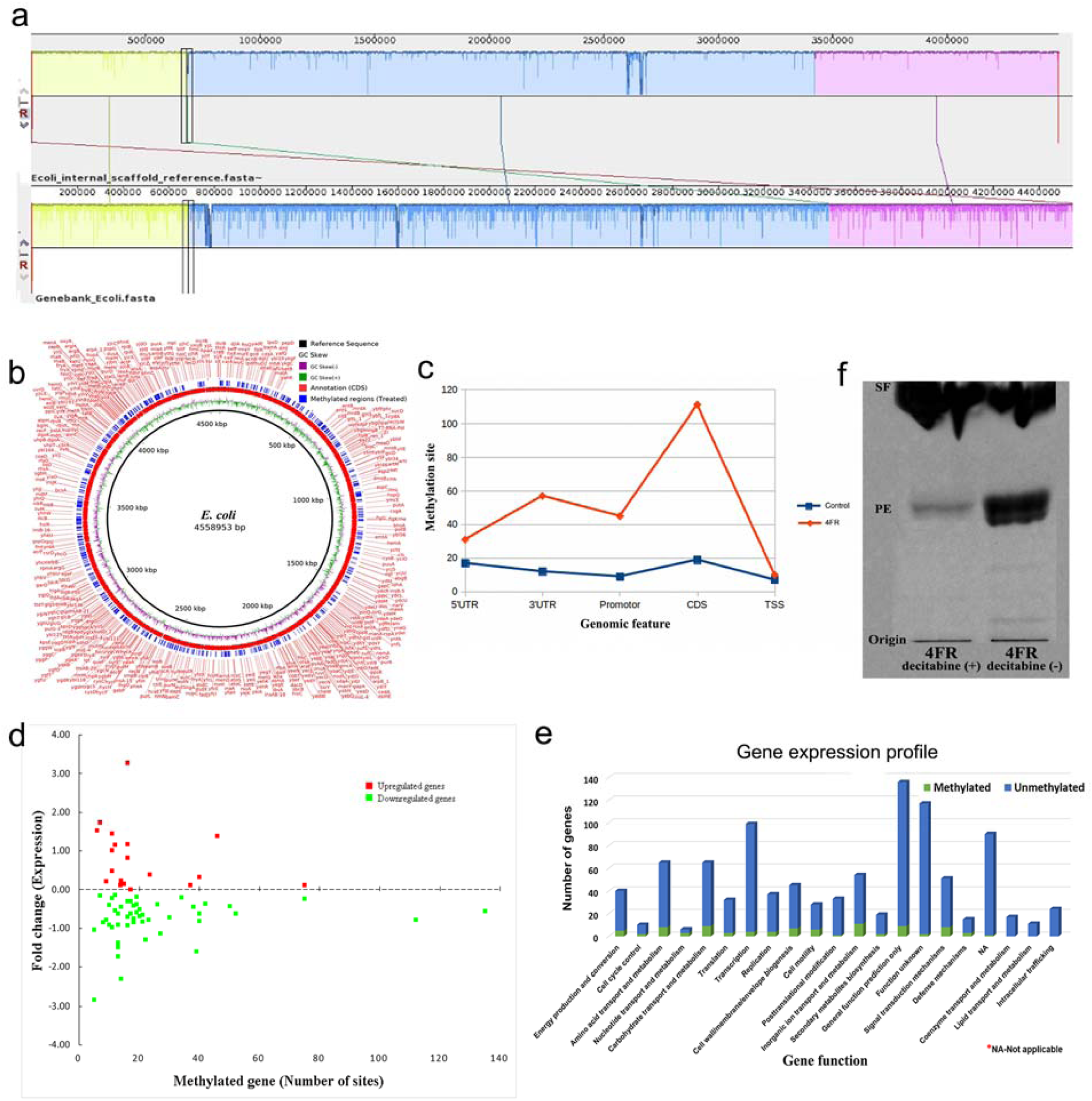
Comparison of Assembled Scaffold: Sequence similarity search of two strains 4FR and control. Reads were processed by poretools for converting fast5 files to Fasta format. Assembly was carried out using OLC based *denovo* assembler such as CANU, miniasm, and NaS. The assembler takes reads and assembles them into contigs using overlapping consensus information. Reads were error corrected, trimmed and assembled into a single scaffold. Sequence similarity search between assembled *E.coli* and Genbank *E.coli* Accession No. CP001509.3 showed that 98% identity at an E-value cut-off of e-5 (<0.00001). We have sequenced 239.31 and 724.32 Million bases (1D read and 2D read) for control and 4FR samples respectively (Extended Data Table S3a, b). Sequence similarity search of assembled *E.coli* showed that 98% identity at an E-value cut-off of e-5 (<0.00001) and have predicted 3715 genes from an assembled scaffold and 97% of the genes were annotated (Extended Data Table S3c). **4b.** Methylation sites (i.e., a locus where methylation occurred) of the control sample and 4FR sample represented in circos plot using BLAST Ring Image Generator (BRIG) programme. BRIG generated circos plot that showed a comparison of control and 4FR--methylation sites on the genome. *Tracks from outside to inside are as follows*: The plot summarizes the number of genes methylated in the 4FR (blue). The CDS of the gene on the strand is showed in the red. The histogram shows the GC Skew (green and purple) of 4FR genome, where GC Skew (G-C) is marked in purple (-) and GC Skew (G+C) in green (+). The black circle inside the circos plot is the reference sequence (control), which we used to compare the 4FR strain. The analysis section of the genome is mentioned in the materials and method. Nabil-Fareed Alikhan, Nicola K Petty, Nouri L Ben Zakour and Scott A Beatson BLAST Ring Image Generator (BRIG): simple prokaryote genome comparisons BMC Genomics 2011 12:402. **4c.** Global methylation in the genome: The genomic features (UTRs, Promotor, CDS, and TSS) of methylated regions from the control (blue) and 4FR (red) has been identified and plotted. A large number of CDS methylated in 4FR comparatively control. Followed by methylation in UTR, promotor, and TSS is enriched in the 4FR. Identified genes were differentially expressed in 4FR classified according to its function represented in X-axis. A total number of genes involved is shown in Y-axis. This obtained data methylated and unmethylated genes were compared with microarray (transcriptome) data. The number of genes identified as unmethylated is denoted in blue and the methylated genes in green. We reported methylation sites in genomic feature of control and 4FR. Gene expression probe used in the gene expression array was mapped to the assembled and used to map the individual gene’s expression status to its overall methylation count. We observed that the genes were distinctly correlating with down-regulated genes. Unmethylated genes grouped with the up-regulated genes. Methylation in genomic regions with a score p >0.99 was aggregated as counts and was correlated with gene expression data for differential p=<0.05. **4d.** Correlation of methylation genes effects to the unmethylated genes expression: Control versus 4FR was computed for higher posterior probability (>0.99) of the methylated sites. Gene-expression probe used in the gene expression array was mapped to the assembled and used to map each gene’s expression status to its overall methylation count. We observed that methylated genes were distinctly correlating with down-regulated genes. Unmethylated genes grouping with up-regulated genes (Methylation in genomic regions with a score >0.99 were aggregated as counts and were correlated with gene expression data for differential p=<0.05). 4e. Graphical representation of the methylated gene expression profile of upregulated and downregulated genes of 4FR. **4f.** One-dimensional-TLC analysis for Phosphatidylethanolamine content in 4FR with demethylation drug decitabine, the cells were harvested, and extracted lipids were resolved on a TLC plate. PE, phosphatidylethanolamine; SF, solvent front. The lipids were separated by using the solvent system chloroform: methanol: ammonia (65:25:5, v/v) as the first dimension. Showing the effect of decitabine on PE synthesis in 4FR with (+) and without (-) decitabine.

Upon analyzing the genome-wide methylation profile, we found that there was global methylation in the 4FR when compared with the control (Fig. 4b). It was observed that the methylation did not follow a pattern and the various regions of the genome seem to be randomly methylated (Fig. 4c). These grounds change in the gene expression in 4FR as compared to control. It is being depicted in Fig.4d showing microarray transcriptome data (GEO accession ID SO_5594: GSE87438) comparing with 68 methylated genes. A range of 5 to 135 sites are methylated in individual genes, and this affects the gene expression up (18 genes) and down (50 genes). Also, this effect extends a change in unmethylated genes expression level as compared to control. Out of the 1738 unmethylated genes, 852 genes are up-regulated, whereas the remaining is downregulated (Fig. 4e, Extended Data Table S4c). Similarly, a few cytosine methylations in the *E. coli* genome affected about 510 genes expression level(*19*). The cytosine methylation could alter the mechanical properties of DNA, thereby affecting the regulation of gene expression(*20*). The differences in gene expression, which has caused the change in metabolic pathways and cell envelope thickening could be due to the global methylation of genes observed in the 4FR. The use of decitabine, methylase inhibitor, in the 4FR culture reduced synthesis of PE to 6.7 times as compared to 4FR without decitabine (Fig. 4f), indicating that methylation is responsible for the cause of PE synthesis in 4FR.

The antibiotic stress on bacteria makes it struggle for its survival. In our study, when an opportunity to survive was presented in the form of an antibiotic resistance gene, the organism has adapted itself for the recombination to take place even in the absence of RecA gene. It was possible due to the global methylation as a defensive mechanism that occurred in the genome of 4FR, which caused the change in gene expression, protein, profiles, lipid synthesis that are being observed in our study. These changes lead to the differential regulation of molecular pathways and cell envelope thickening, which could allow the recombination to take place. The epigenetic mediated phenotypic alternation that took place in the organism helped in providing the window required for recombination of four fragments. This could, in future, be used as a model to study the adaptations of pathogenic organisms, thereby combating antibiotic microbial resistance (AMR). This phenomenon could be applicable to any form of life or life-threatening situation which leads us to the belief that the well-characterized pathway may not be the solicit ones to carry out the functions and there may be other pathways still unexplored.

## Acknowledgments

We would like to thank Dr. Renu Paricha, Head of TEM facility, National Center for Biological Sciences (NCBS) for providing training in Electron Microscopy (TEM). We would also like to thank Dr. Vikas Kumar and Mr. Prashanth K of the Mass spectrometry facility in CCAMP (Centre for cellular and molecular platforms) for providing service in generating LC-MS Data of label-free proteomics analysis. Also, we thank Dr. Kannan C-CAMP for providing service measuring metabolites by UHPLC-MS/SRM method. We also thank for Genotypic Technology, Bangalore India for providing service for Microarray, Nano-pore and methylation services and data analysis. We author thank Mr. Lakshmi Narayanan Gopalan, Soumya Kariyadan Veetil, Narendrakumar Sekar, and Rohini Srivastava for their help.

## Author Contributions

MN conceptualized the project. SM, SB, and VV performed the experiments. RR and VV designed lipidomics experiments. MN wrote the manuscript. MN, SM, SB, CC, RR, and VV proofread and verified the data of the manuscript. SM, SB, and VV made the figures.

## Supplementary Materials

### Materials and Methods

#### Preparation of DNA fragments for recombination

The DNA molecules used in this study were derived from various sources, as described in Supplementary Table S2. All strains containing plasmids were propagated in LB medium containing appropriate antibiotic and plasmid isolated using Sigma’s GenElute™ plasmid miniprep kit and eluted in TE buffer. The resulting plasmids were used as templates for PCR using Phusion DNA polymerase with HF buffer (NEB). For Phusion polymerase reactions, a final concentration of 1x HF buffer, 200 µM dNTPs, 0.5µM of forward and reverse primer each (Supplementary Table S2), 3% DMSO, 10ng of DNA, 0.5 units of Phusion DNA polymerase were used and made up to 25 µl with ultra-pure water from GIBCO. Further, the PCR products were treated with DpnI to eliminate template contamination and gel extracted using Sigma’s GenElute™ Gel Extraction kit. A second identical PCR was performed to generate sufficient amplicons and salts were removed using Sigma’s GenElute™ PCR purification kit. The final product was eluted in water for cloning purposes. Antibiotic concentrations, primers, and overhangs are described in Supplementary Table S2. All fragments were tested to be mother template free by transforming them individually, which did not result in colonies.

#### Transformation of linear DNA fragments

Transformation was performed both by heat-shock and electroporation. 40 µl of competent cells (Supplementary Table S1) were thawed on ice, and equimolar concentration of linear DNA fragments with appropriate overhangs as described in supplementary table S1 was added. After incubation for 30 minutes, the cells were heat-shocked for 45 seconds at 42°C. This was followed by 2-minute incubation and 1ml SOC, or LB was added. Further, incubation was carried out in a 37°C incubator at 200 rpm for 90 minutes. The cells were pelleted at 6000 rpm for 20 seconds and plated on appropriate antibiotics and incubated at 37°C overnight. Transformants were screened by colony PCR and confirmed by sequencing.

Electroporation was performed in 2mm cuvettes from Bio-Rad in the Bio-Rad GenePulser. 40 µl of cells were transformed with an equimolar concentration of DNA fragments and pulsed at 3000 kV with a capacitance of 2000 F. A time constant of 5 was maintained for all electroporation. Cells were incubated at 37°C overnight and transformants were confirmed by sequencing.

#### Adaptive test 1

Two identical 4 fragment transformations were carried out in BLR (DE3) and pooled. Half was plated on LB agar containing Kanamycin (LB-K), while the other was serially diluted in a range of 10^−1^-10^−6^ in 100 µl SOC, resuspended and plated on plain LB plates (LB-P). After incubation at 37°C overnight, 1ml of LB was used to ensure all colonies from LB-K and 10^−1^, 10^−2^, 10^−3^ dilutions of LB-P were completely scooped from plates and inoculated into 100ml of LB containing kanamycin. Cultures were grown at 37°C at 200 rpm and monitored the growth at OD600. A graph was plotted against OD600 versus time. To further validate LB-K, LB-P colonies were scrapped and plated on LB-K. The above experiment was performed in triplicates (Extended Data Fig. S1).

#### Adaptive test 2

Two-identical 4 fragment transformations were performed and pooled. Further, half were plated on LB-K and others on LB-P, followed by incubation overnight at 37°C. All resulting colonies were scraped and re-plated on LB-K plates. The above was performed in triplicates (Extended Data Fig. S1).

#### Adaptive Window

4FR clones were plated on LB agar plates of kanamycin concentrations ranging from 10-60μg/ml, with untransformed BLR and BLR containing pET28a as controls. The above was performed in triplicates. Colonies were counted after 36 hrs and incubation at 37°C (Fig. 1c).

#### Block Preparation protocol for TEM (FEI Tecnai™ G2 F20)

Cells were grown till exponential phase from overnight culture and media was separated from the cell pellet by centrifugation and resuspended in 4% glutaraldehyde in 0.1M sodium cacodylate buffer (pH 7.4). This was stored at 4°C overnight. Next, the fixative was removed by centrifugation at 3000 rpm at 4° C for 15 minutes. The supernatant was discarded and further resuspended in fresh 0.1 M sodium cacodylate buffer pH 7.4. This was repeated 3 times at 3000 rpm for 15 mins at room temperature (RT) until the fumes of glutaraldehyde were gone. Further, the supernatant was discarded and the pellet was resuspended in 1% osmium tetraoxide (aqueous) at RT for 75 mins. Black pellet color was observed confirming the fixing. The pellet was washed with 0.1M sodium cacodylate buffer pH 7.4 until the fumes of osmium tetraoxide vanished. The supernatant was then replaced with 70% and 80% ethanol and centrifuged for 1 hr at room temperature each and supernatant discarded. Following this pellet was resuspended in 2% Uranyl acetate, which was prepared in absolute ethanol and light protected. This was incubated in dark for 1 hr at room temperature and centrifuged to remove supernatant. Absolute ethanol was added into the pellet and incubated for 1 hr at room temperature and centrifuged to remove supernatant. Next, the supernatant was replaced with propylene oxide twice each for 15 mins at room temperature. The pellet was incubated in 1:1 ratio of propylene oxide: resin mixture which was freshly prepared. Further, it was rotated in a rotator overnight. Propylene oxide resin mixture was removed carefully and replaced with only resin and kept on a rotator for 2 hrs and 4 hrs. At the end of 4 hrs of rotation in the resin, the fresh resin was poured into the mold, and the pellet was picked and poured into the mold. Mould was kept in a hot air oven at 60°C for 72 hrs (or until the blocks are hard enough to come out of molds easily). The blocks were trimmed using razor blades to remove most of the resin from the sample cutting edge, making it look like a pyramid. Then the thin sections were checked for dissection on the Ultra-microtome by picking thick (approximately 200-400 nm) sections on a drop of water on a glass slide. This slide was placed on a hot plate until the water evaporated completely. The sections were stained with Toluidine blue stain for 20 seconds and washed with water and dried on a hot plate to check quality. After toluidine blue staining, thinner (50-100 nm) sections were cut and picked onto uncoated 400 square mesh grids. Grids with sections were stored at 60°C for 3-4 hrs in a hot air oven.

#### Staining protocol for imaging in TEM

The grid containing sections was stained with 2% Uranyl acetate (aqueous) for 1 hr. The grids were washed thoroughly in a beaker containing distilled water by dipping for approximately 30 seconds in each beaker of water, excess water was removed with filter paper. The grids were then stained with 0.04% Lead citrate solution for 5mins in the presence of Sodium hydroxide pellets. Next, the grids were washed thoroughly by dipping in a beaker containing distilled water for 30 seconds twice and excess water removed with filter paper. Dry grids with sections were then kept the side up under table lamp for 2 hrs and Imaged in TEM.

#### Lipids extraction for thin layer chromatography (1D and 2D) and lipidome analysis

A single colony of *E. coli* cells was grown at 37 °C, overnight in Luria-Bertani (LB) media. The cells were harvested by centrifugation, and the cell pellet was washed thrice with water. An equal amount of cells (A_600_= 5) was used for lipid extraction. For demethylation study, 5µg of decitabine was added into the media. The total lipids were extracted by 1:1:1 (v/v) chloroform: methanol: 2% orthophosphoric acid. The extracted lipids were separated by two-dimensional silica-TLC using chloroform: methanol: ammonia (65:25:5, v/v) as the first dimension and chloroform: methanol: acetone: acetic acid: water (50:10:20:15:5, v/v) as the second dimension. The intensity of the lipid spots was quantified by ImageJ software. The values are represented as the mean (±SD) of three independent experiments performed in triplicates. Significance was determined at ***p< 0.001.

For lipidome analysis, A_600_= 5.0 OD cells were harvested, for lipid extraction. The dried lipids were dissolved in 1:2 ratio of chloroform: methanol containing 10 mM ammonium acetate. The samples were injected to AB Sciex TripleTOF™ 5600 System using MS/MS ALL method(*21*). The molecular species of PE were identified in positive mode. The lipids were analyzed by Peak View and Lipid View software (AB SCIEX). The amount of phosphatidylethanolamine (PE) was represented as a fold change of total lipids. Data are mean of three independent experiments with ±SD performed in triplicates.

#### Label-Free LC-MS Mass-spectrometry and Data-Analysis

*E.coli* cells (Control and 4FR clones) were grown till late exponential phase from overnight culture and lysed by sonication. This was centrifuged at 13,000 rpm for 30 minutes at 4°C to remove media and cell debris. An equal amount of samples (50 µg) were used for processing by acetone precipitation. For reduction, 10mM Dithiothreitol (DTT) was added and kept at 56°C for 30 min. For alkylation, 50mM iodoacetamide (IAM) was kept at room temperature in dark for 20 min. For digestion, sequencing grade Typsin in protein: substrate ratio of 1:25 was used. Next, overnight digestion was performed at 37°C. The reaction was then stopped by adding concentrated formic acid. After digestion, tryptic peptides were vacuum dried and re-dissolved in 50 microliters of 2% acetonitrile with 0.1% formic acid. Out of 50, 1 µl was injected into the mass spec for data acquisition. Data was acquired using Thermoscientific LTQ Orbitrap Discovery. Further, proteomic data was filtered and analyzed in Progenesis QI blasted across Swiss-Prot and RefSeq databases at a p-value of 0.05%.

#### Microarray and data analysis

##### RNA Extraction and RNA Quality Control

*E. coli* pellet was re-suspended in 300 µl of 5 mg/ml lysozyme and incubated at room temperature (RT) for 30 min. Isolation of RNA from *E. coli* was carried out by using Qiagen RNeasy mini kit as per Manufacturer’s guidelines including DNase treatment step. The purity of the RNA was assessed by using the Nanodrop Spectrophotometer (Thermo Scientific; ND-2000) and the integrity of the RNA was analyzed on the Bioanalyzer (Agilent; 2100). We considered RNA to be of good quality based on the 260/280 values (Nanodrop), rRNA 28S/18S ratios and RNA integrity number (RIN) (Bioanalyzer). The sample labeling was performed using Quick-Amp Labeling Kit, One Color (Agilent Technologies, Part Number: 5190-0442). Approximate amount of RNA from each of the sample was denatured along with WT primer along and T7 polymerase promoter primer. The cDNA master mix was added to the denatured RNA sample and incubated at 40°C for 2 hrs for double-stranded cDNA synthesis. Synthesized double-stranded cDNA was used as a template for cRNA generation. cRNA was generated by *in vitro* transcription and the dye Cy3 CTP incorporated during this step and incubated at 40°C for 2:30 hrs. The Cyanine 3-CTP labeled cRNA sample was purified using a Qiagen RNeasy column (Qiagen, Cat No: 74106). The concentration of cRNA and dye incorporation was determined using Nanodrop-1000.

About 4 micrograms of labeled Cyanine 3-CTP cRNAs were fragmented at 60°C for 30 minutes and the reaction was stopped by adding 2X GE HI-RPM hybridization buffer (Agilent Technologies, In situ Hybridization kit, Part Number5190-0404). The hybridization was carried out in Agilent’s Surehyb Chambers at 65°C for 16 hrs. The hybridized slides were washed using Gene Expression Wash Buffer1 (Agilent Technologies, Part Number 5188-5325) and Gene Expression Wash Buffer 2 (Agilent Technologies, Part Number 5188-5326) and were scanned using Agilent Scanner (Agilent Technologies, Part Number G2600D). Data extraction from Images was done using Feature Extraction software Version Feature. Extracted raw data was analyzed using Agilent GeneSpring GX software. Normalization of the data was performed in GeneSpring GX using the 75th percentile shift method [Percentile shift normalization is a global normalization, where the locations of all the spot intensities in an array are adjusted. This normalization takes each column in an experiment independently, and computes the n^th^ percentile of the expression values for this array, across all spots (where n has a range from 0-100 and n=75 is the median. It subtracts this value from the expression value of each entity] and fold change values were obtained by comparing test samples with respect to specific control samples. Significant genes up-regulated fold > 1 (logbase2) and down-regulated < -1 (logbase2) in the test samples with respect to control sample were identified. Statistical student T-test p-value among the replicates was calculated based on Volcano plot algorithm. Differentially regulated genes were clustered using hierarchical clustering based on Pearson coefficient correlation algorithm to identify significant gene expression patterns. Genes were classified based on functional category and pathways using Biological Analysis tool DAVID (http://david.abcc.ncifcrf.gov/).

#### Library preparation and third-generation sequencing

Control and 4FR DNA was run on a 0.8% agarose gel for integrity check. Followed by Genomic DNA (2000ng) was fragmented (Covaris, Inc., MA, USA) in a water bath maintained at 4°C. A sheared DNA sample was purified using 1x AmPure beads (Beckman Coulter life sciences, IN, USA). End-repair was performed using NEBnext ultra II end repair kit (New England Biolabs, MA, USA). End-repaired DNA was cleaned up with 1x AmPure beads. Adapter ligations (HPA and then HPT) were performed for 15 minutes each using NEB blunt/ TA ligase (New England Biolabs, MA, USA). Library mix was cleaned up using 50 µl of myOne C1 beads and finally eluted in 25 µl of elution buffer. ATEL∼170.6 ng (7.08 ng/ µl) of yield was obtained and used for sequencing. Sequencing was performed on MinION Mk1 (Oxford Nanopore Technologies, Oxford, UK) using flowcell ID FAD03118 in a 48 hr sequencing protocol. Oxford Nanopore passes an ionic current through nanopores and measures the changes in current as biological molecules pass through the nanopore. The ‘bulk data’ are segmented into discrete ‘events’ of similar consecutive measurements. The 5-mer corresponding to each event is inferred using a statistical model. We have sequenced 239.31 and 724.32 Million bases (1D read and 2D read) for control and 4FR samples respectively.

#### Detection and mapping of methylation in genome data

Reads were processed by poretools for converting fast5 files to Fasta format. Assembly was carried out using OLC based *de-novo* assembler such as CANU, miniasm, and NaS. The assembler takes reads and assembles them into contigs using overlapping consensus information. Reads were error corrected, trimmed and assembled into a single scaffold. Sequence similarity search between assembled *E.coli* and Genbank *E.coli* Accession No. CP001509.3 showed that 98% identity at an E-value cut-off of e-5 (<0.00001). The coding sequence of the assembled scaffold sequence was predicted using RAST (Rapid Annotation Using Subsystem Technology) programme^5^. We have predicted 13249 CDS, 249 tRNA and 66 rRNA from assembled scaffold *E.coli* sequence. Assembled *E.coli* scaffold sequence used as a reference for reading mapping and methylation calling. Ionic current events from basecalled Minion reads (.fast5 files) were mapped to the reference sequence using SignalAlign programme. SignalAlign programme supported aligning multiple cytosine variants (methylation variants) to a position. We separately aligned the events from the complement and template strand to the reference sequence with model and marginalize over the HMM’s states to obtain the posterior probability on the methylation status.

#### Identifying methylation in genomic features

We have identified global methylation in genomic feature (UTRs, Promotor, CDS, and TSS) from control and 4FR samples. A total of 111 CDS methylated in 4FRsample and 19 CDS in the control sample. Followed by methylation in UTR, promotor and TSS regions were analyzed in control and 4FR sample. We investigated whether the newly methylated regions localized to specific gene features. Regions of methylation were highly associated with genes in 4FRsample. 4FRhad higher than expected methylation enrichment within 0.5–1 kb upstream or downstream of genes and promoter region.

RecA gene retrieved from genebank database and mapped back to the scaffold. Mapping result showed RecA absence in the sample, and also we have checked tetracycline resistance gene methylation in 4FR sample and control sample. Methylation obtained in the tetracycline resistance gene in 4FR sample but not in the control sample.

Followed by the methylation site in the hotspot region was carried out, we have found four genes that were methylated in 4FR sample. The above evidence strongly showed the methylation occurred prominently in 4FR sample.

## Supplementary Tables

**Supplementary Table S1.** Genotypic and source of the *E. coli* strain used in the study.

| Strain | Genotype | Source |
| --- | --- | --- |
| *E. coli* (BLR) | *F<sup>-</sup> ompT hsdS<sub>B</sub> (r<sub>B</sub><sup>-</sup> m<sub>B</sub><sup>-</sup>) gal lac ile dcm Δ(srl-RecA)306::Tn10 (Tet<sup>R</sup>)* | Novagen |

**Supplementary Table S2.**
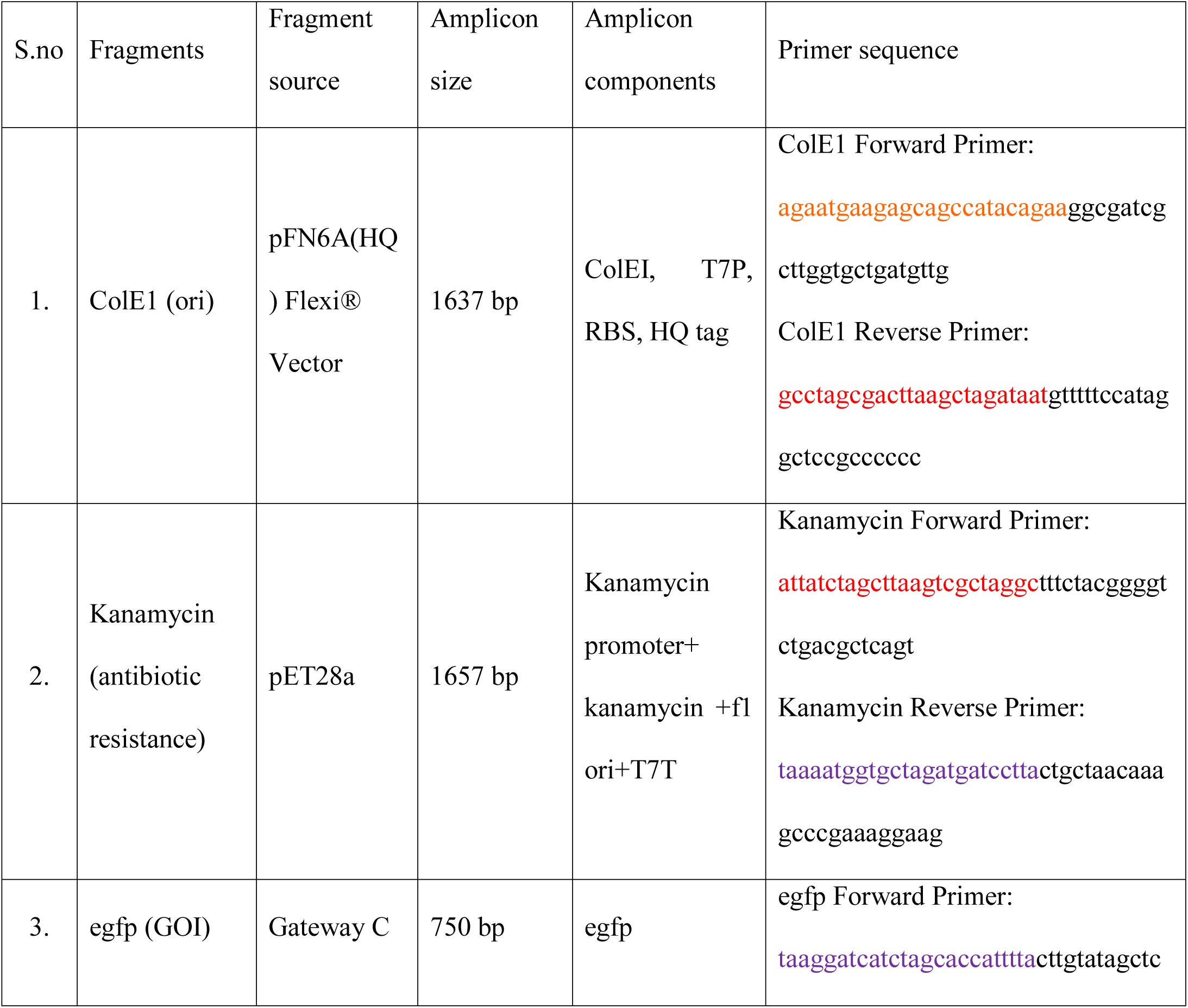
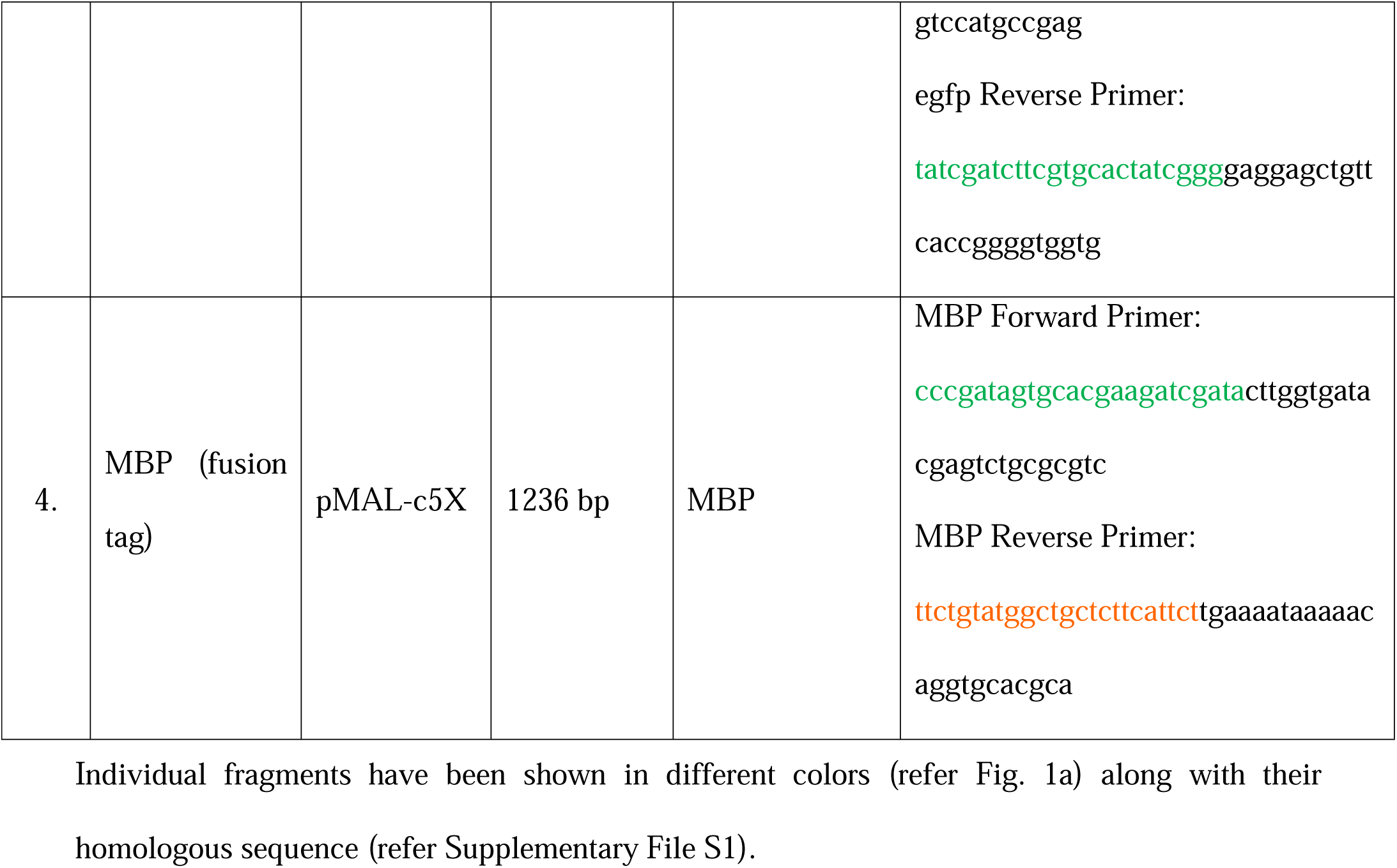
Primer sequence and amplicon size of four fragments.

## Supplementary File S1

### 4FR sequence

Individual fragments has been shown in different colors (refer Fig. 1a) along with their homologous sequence (refer Supplementary Table S2).

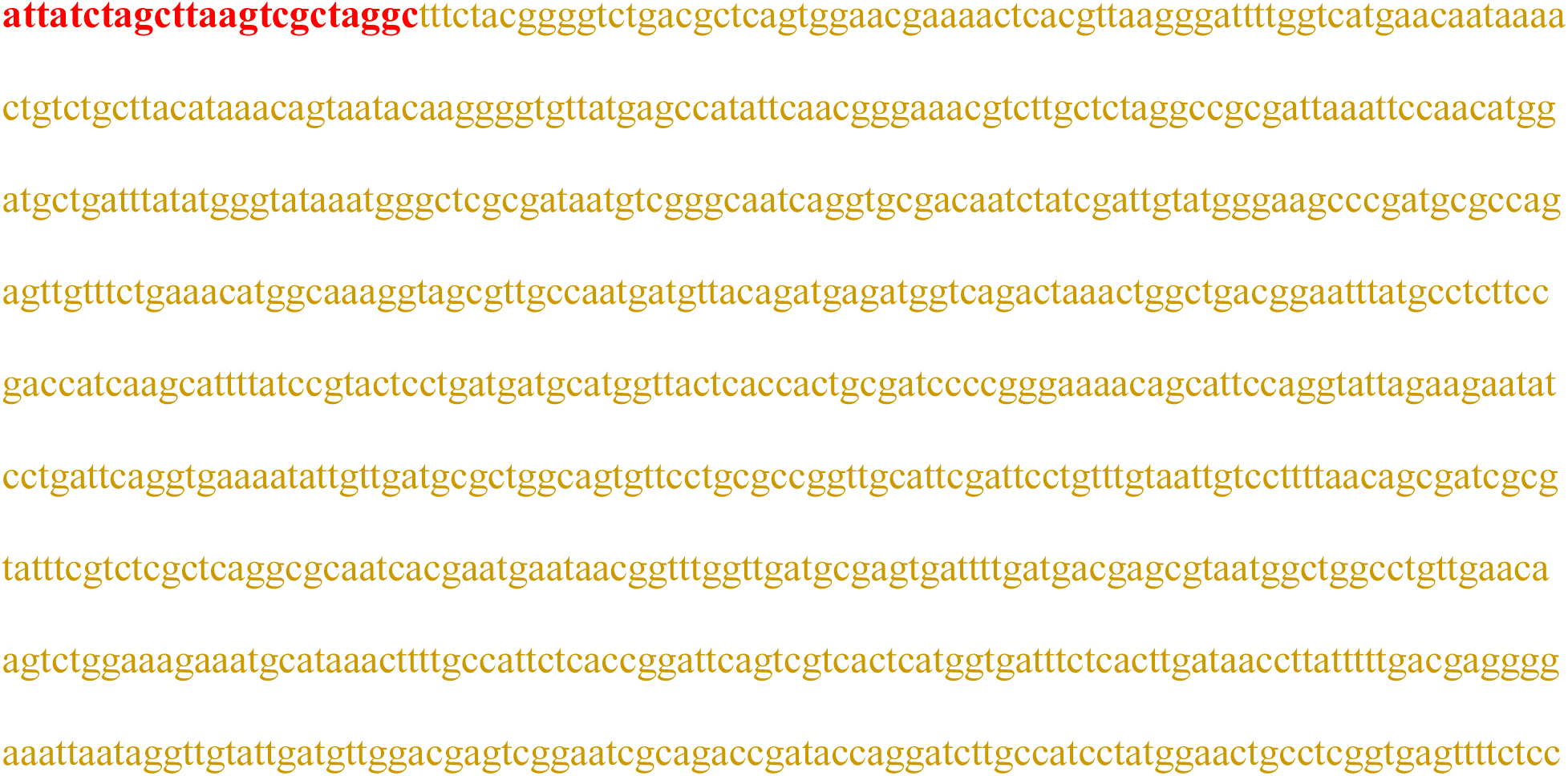

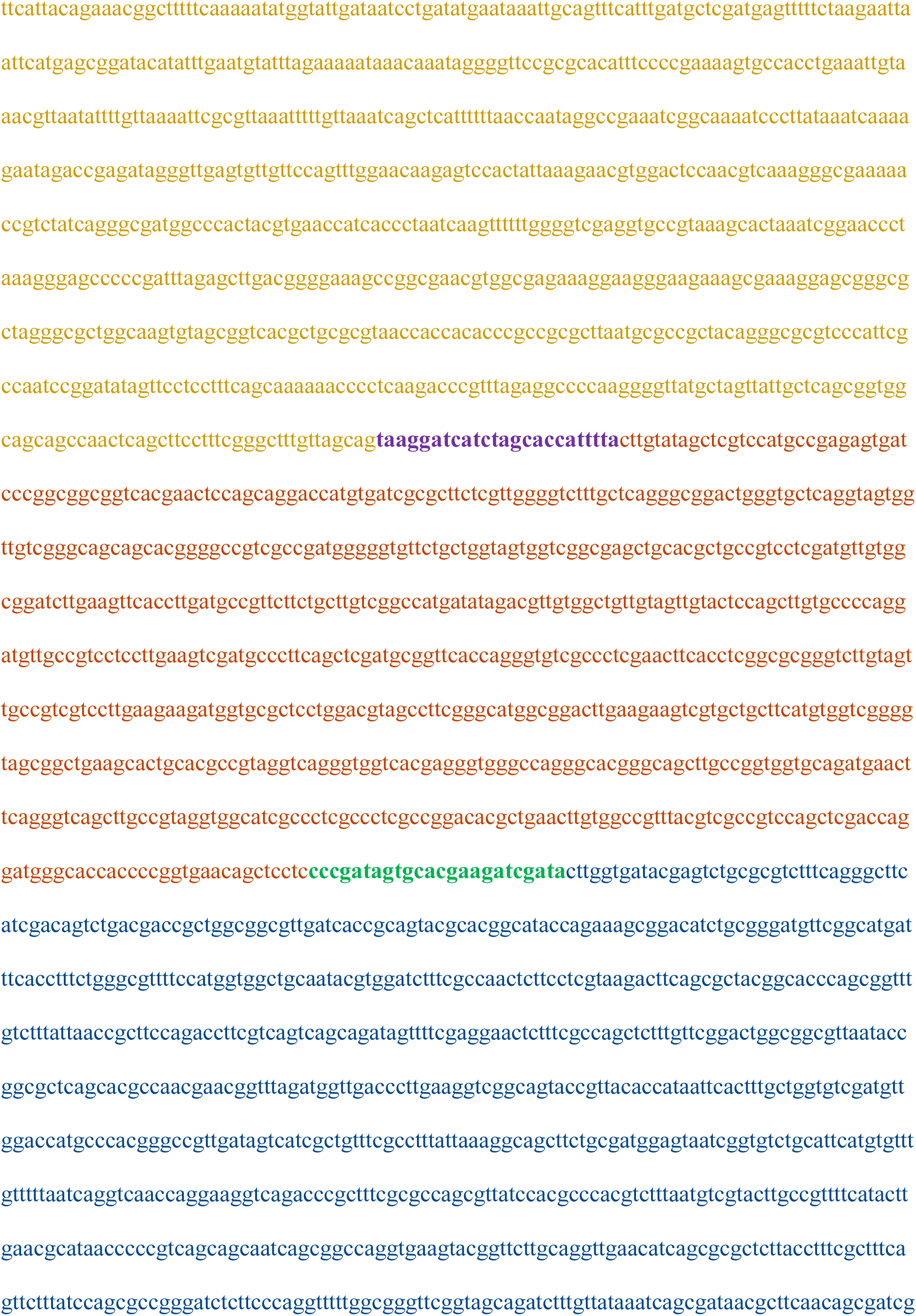

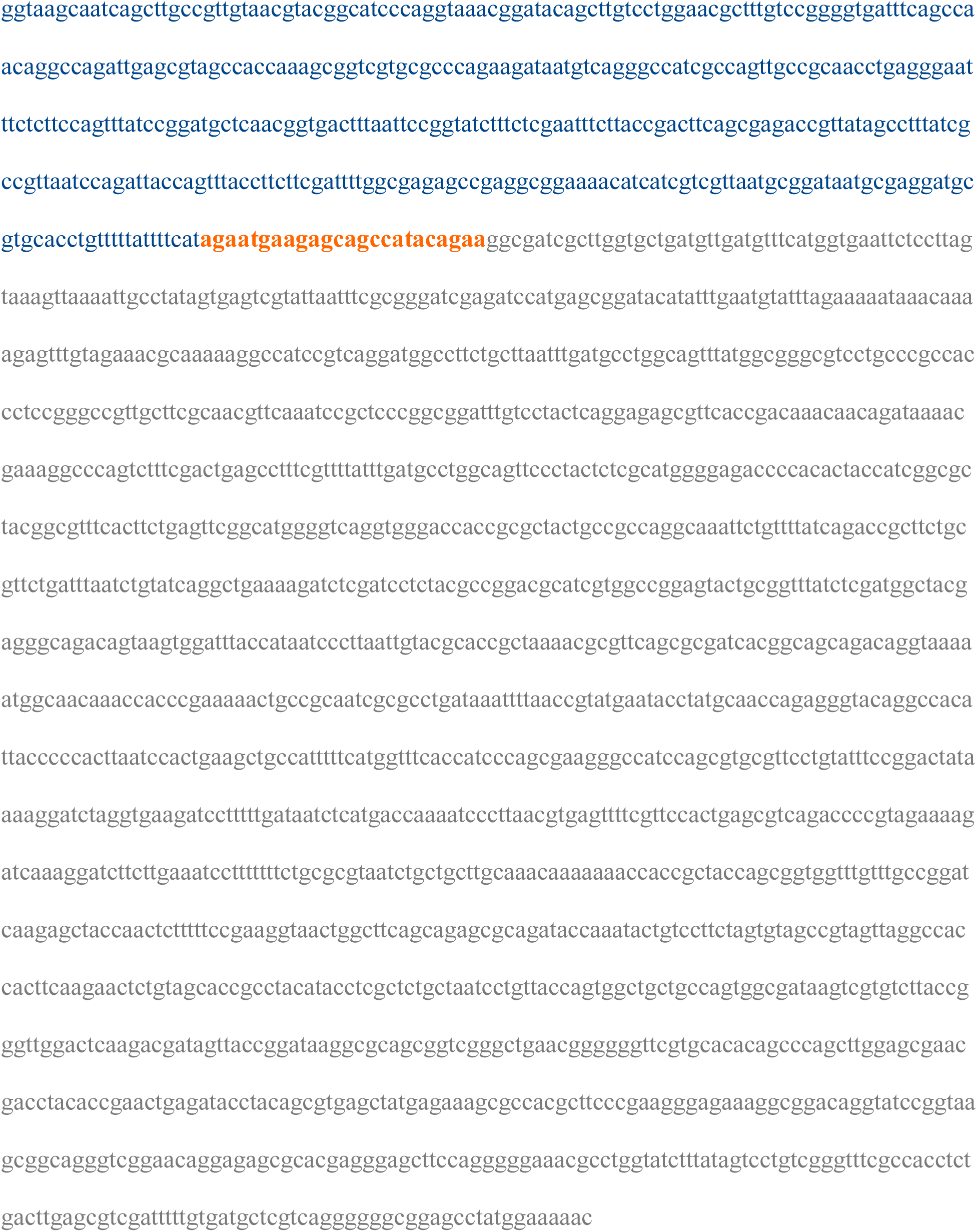

## Extended Data Figure Legends

Extended Data Fig. S1. **RecA independent homologous recombination by stress a.** Transformation of Non-DpnI treated and DpnI treated shows 4FR occurs only by stress-mediated RecA independent homologous recombination. If a circular plasmid bearing similar resistance is introduced along with the 4FR fragments, 4FR recombination fails. **b.** Illustration describing the transformation of two identical linear fragment(s). 4FR recombination was obtained only upon exposure to stress by LB-kanamycin antibiotic (LB-K) at 30 µg/ml kanamycin and failed to occur in the no-stress LB-plain (LB-P) plate, and it showed subsequent growth in antibiotic-containing LB-broth. **c.** Left: Amplification of eGFP (∼700bp) from Non-DpnI colonies, Lane C is control amplification for eGFP, Lane M is 1Kb plus DNA ladder, Lane 1-6 is clones showing no amplification of eGFP; Right: Amplification of eGFP from DpnI treated and gel-extracted C-D colonies, Lane M is 1Kb plus DNA marker, Lanes 1-5 show amplification of eGFP. **c.** Left: Amplification of eGFP (∼700bp) from Non-DpnI colonies, Lane C is control amplification for eGFP, Lane M is 1Kb plus DNA ladder, Lane 1-6 is clones showing no amplification of eGFP; Right: Amplification of eGFP from DpnI treated and gel-extracted C-D colonies, Lane M is 1Kb plus DNA marker, Lanes 1-5 show amplification of eGFP.

