## Supplementary figures and images for "Phenotypic changes of bacteria through opportunity and global methylation leads to antibiotic resistance"

### Extended data figure S1

**Extended data figure S1**

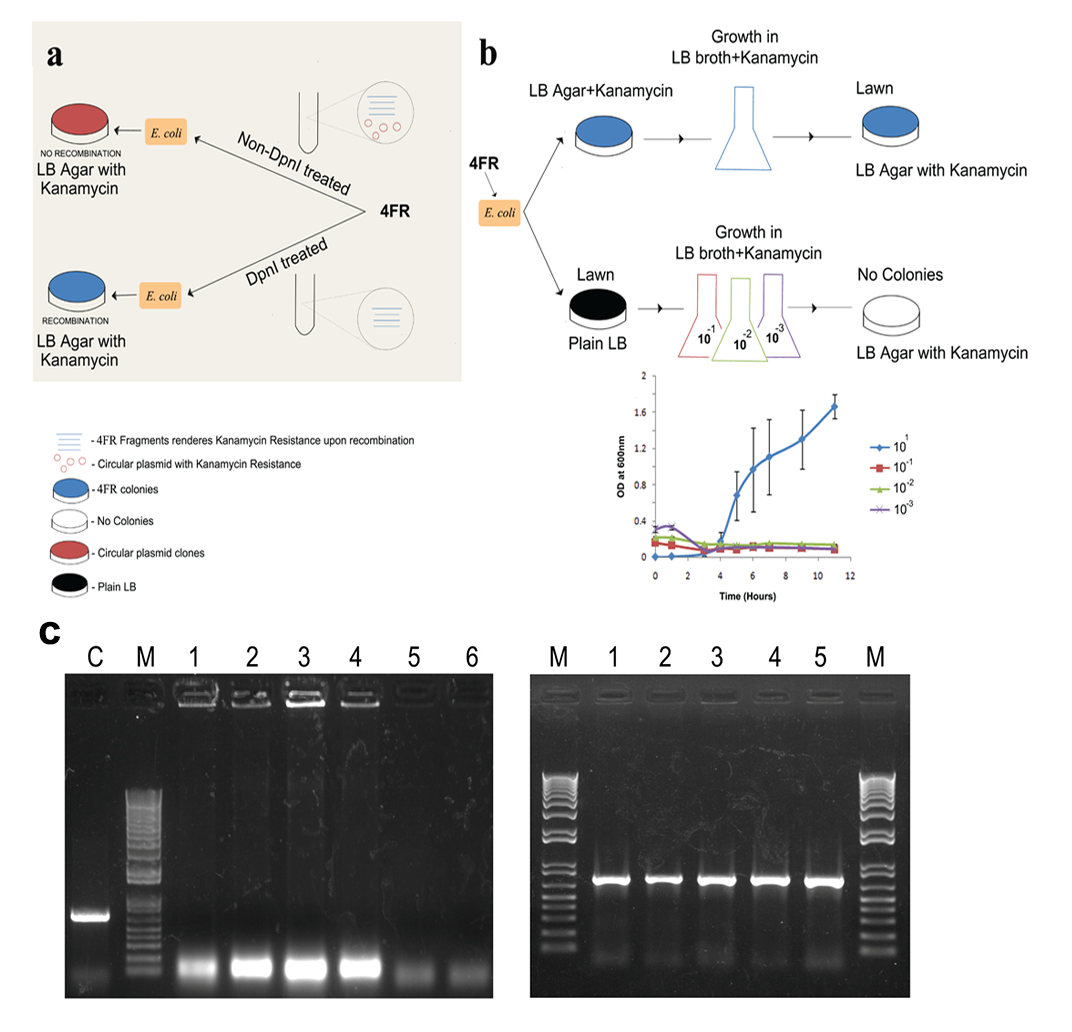
