## Extended data-Table1 for "Phenotypic changes of bacteria through opportunity and global methylation leads to antibiotic resistance"

**Extended Data Table 1**

| Kanamycin (µg/mL) | Control (10^-6^) | 4FR (10^-6^) |
| --- | --- | --- |
| 250 | 84 ±4 | 254 ±7 |
| 500 | 12 ±1 | 157 ±8 |
| 1000 | 0 | 104 ±7 |
| 2000 | 0 | 47 ±3 |
| 4000 | 0 | 13 ±2 |

Single colony of control and 4FR were grown at 50 µg/mL kanamycin. Cells were taken at OD_600nm_ 0.6 and serial diluted up to 10^-6^ and plated on different concentration of kanamycin. The number of colonies obtained are represented as the mean (±SD) of the triplicates. The control was transformed with CkeM plasmid that was isolated from the 4FR.
