## Extended data-Table3a, b, c for "Phenotypic changes of bacteria through opportunity and global methylation leads to antibiotic resistance"

**Extended Data Table 3a.**

| ***E.coli* Control and 4FR 1D reads statistics** | | |
| --- | --- | --- |
|  | **Control** | **4FR** |
| Reads Generated | 60159 | 83055 |
| Maximum read Length | 659377 | 274706 |
| Minimum read Length | 1000 | 1000 |
| Median read Length | 2489 | 14178 |
| Total reads Length | 208854672 | 589841476 |
| reads >= 100 bp | 60159 | 83055 |
| reads >= 200 bp | 60159 | 83055 |
| reads >= 500 bp | 60159 | 83055 |
| reads >= 1 Kbp | 60159 | 83055 |
| reads >= 10 Kbp | 2125 | 14370 |
| X-Coverage | 46.55 | 131.46 |

**Extended Data Table 3b.**

| ***E.coli* Control and 4FR 2D reads statistics** | | |
| --- | --- | --- |
|  | **Control** | **4FR** |
| Reads Generated | 11367 | 31336 |
| Maximum read Length | 22830 | 57595 |
| Minimum read Length | 1000 | 1000 |
| Median read Length | 1419 | 3789 |
| Total reads Length | 30461354 | 134486615 |
| reads >= 100 bp | 11367 | 31336 |
| reads >= 200 bp | 11367 | 31336 |
| reads >= 500 bp | 11367 | 31336 |
| reads >= 1 Kbp | 11367 | 31336 |
| reads >= 10 Kbp | 43 | 1561 |
| X-Coverage | 6.79 | 29.97 |

**Extended Data Table 3c. Summary of the Gene annotation**

| Total no of Predicted Genes | 3715 |
| --- | --- |
| Total no of annotated Genes | 3640 |
| % of Annotated genes | 97 |
| Total no of genes methylated | 111 |
