## Extended data-Table 4a & 4b for "Phenotypic changes of bacteria through opportunity and global methylation leads to antibiotic resistance"

| **Gene Name** | **Number of sites methylated** |
| --- | --- |
| Transcriptional repressor of the lac operon | 24 |
| Helicase PriA essential for oriC/DnaA-independent DNA replication | 44 |
| Maltodextrin phosphorylase | 54 |
| Xylulose kinase | 12 |

**Extended Data Table 4a:** Methylation in the hot spot genes in *E.coli* genome (4FR)

**Extended Data Table 4b:** Number of sites methylated in 4FR with tetracycline resistant gene

| Organism | Feature | Start-Coordinate | End-Coordinate | Strand | Gene-Name | Number of sites methylated in 4FR |
| --- | --- | --- | --- | --- | --- | --- |
| *E.coli* | CDS | 2607384 | 2608506 | + | Tetracycline efflux protein TetA | 43 |
